# A bacterium-like particle vaccine displaying Zika virus prM-E induces systemic immune responses in mice

**DOI:** 10.1101/2021.08.13.456215

**Authors:** Hongli Jin, Yujie Bai, Jianzhong Wang, Cuicui Jiao, Di Liu, Mengyao Zhang, Entao Li, Pei Huang, Zhiyuan Gong, Yumeng Song, Shengnan Xu, Na Feng, Yongkun Zhao, Tiecheng Wang, Nan Li, Yuwei Gao, Songtao Yang, Xianzhu Xia, Hualei Wang

**Affiliations:** Key Laboratory of Zoonosis Research, Ministry of Education, College of Veterinary Medicine, Jilin University, Changchun, China; Changchun Veterinary Research Institute, Chinese Academy of Agricultural Sciences, Changchun, China; College of Veterinary Medicine, Jilin Agricultural University, Changchun, China

## Abstract

The emergence of Zika virus (ZIKV) infection, which is an unexpectedly associated with congenital defects, has prompted the development of safe and effective vaccines. The gram-positive enhancer matrix-protein anchor (GEM-PA) display system has emerged as a versatile and highly effective platform for delivering target proteins for vaccines. In this article, we developed a bacterium-like particle vaccine ZI-Δ-PA-GEM based on the GEM-PA system. The fusion protein ZI-Δ-PA, which contains the prM-E-Δ protein of ZIKV (with a stem-transmembrane region deletion) and the protein anchor PA3, was expressed. The fusion protein was successfully displayed on the GEM surface, forming ZI-Δ-PA-GEM. Moreover, when BALB/c mice were immunized intramuscularly with ZI-Δ-PA-GEM combined with 201 VG and poly(I:C) adjuvants, durable ZIKV-specific IgG and protective neutralizing antibody responses were induced. Potent B cell/DC activation was also be stimulated early after immunization. Remarkably, splenocyte proliferation, the secretion of multiple cytokines, T/B cell activation and central memory T cell responses were elicited. These data indicate that ZI-Δ-PA-GEM is a promising bacterium-like particle vaccine candidate for ZIKV.

**Author summary:** Because Zika virus (ZIKV) infection is considered as an example of “disease X”, the development of a safe and effective ZIKV vaccine is essential. The gram-positive enhancer matrix-protein anchor (GEM-PA) display system has been used in many vaccine studies due to its advantages. In this study, prM-E-△ protein of ZIKV (with a stem-transmembrane region deletion) and the protein anchor PA3 was fusion expressed, termed ZI-△-PA. Then the fusion protein ZI-△-PA could be displayed on the surface of GEM, forming ZI-△-PA-GEM. The author evaluated the immunogenicity of ZI-△-PA-GEM with the 201 VG and poly(I:C) adjuvants. The study demonstrates that ZI-△-PA-GEM induced mice to produce neutralizing antibody and specific cellular immune responses. The author believe that the bacterium-like particle vaccine ZI-△-PA-GEM has the potential to be used as the ZIKV vaccine.

## Introduction

Zika virus (ZIKV), first identified in 1947 from a rhesus monkey in Uganda [1, 2], is a mosquito-borne virus [3] belonging to the genus Flavivirus. In 2015, outbreaks of ZIKV were reported in Brazil, and infection was associated with microcephaly, central nervous system malformations and Guillain–Barré syndrome (GBS) [4]. On 1 February 2016, the World Health Organization (WHO) declared ZIKV a ‘public health emergency of international concern’. Moreover, ZIKV disease was listed as a “Disease X” by the WHO in the “Research and Development (R&D) Blueprint for Action to Prevent Epidemics” [5]. As of July 2019, cases of ZIKV had been reported worldwide, including 87 countries and territories, according to a WHO report [6]. To date, no accepted therapies or vaccines are available for ZIKV, and the WHO has made ZIKV vaccine development a top priority [7]. Substantial progress has been made on developing vaccines, which are essential to control the spread of ZIKV. Moreover, new vaccine search efforts are needed because of the complexity of ZIKV immunity and pathogenesis [8–10]. Neutralizing antibody epitopes exist on the surface of the ZIKV E protein [11, 12], and precursor membrane (prM) protein complexes with the E protein are usually ideal candidates for vaccine development.

The gram-positive enhancer matrix-protein anchor (GEM-PA) is a new bacterial surface display system composed of GEM particles and a protein anchor (PA) [13]. First, the foreign protein and PA are expressed as a fusion. Then, the fusion protein “foreign protein-PA” is displayed on the GEM surface by means of the GEM-PA surface display system [14]. The GEM surface display system has the following advantages. (1) Simple purification of a foreign protein by low-speed centrifugation is convenient and efficient. (2) *Lactococcus lactis* (*L. lactis*) is globally recognized as a safe probiotic [15, 16]. After treatment, GEM has no safety risk as a carrier due to the absence of nucleic acids and proteins. (3) The displayed protein can effectively stimulate a stronger immune response [17]. In this study, the GEM-PA display system was used to display ZIKV prM-E glycoproteins on the GEM surface and study their immunogenicity.

## Materials and Methods

### Viruses, cells and antibodies

The recombinant baculoviruses were grown in *Spodoptera frugiperda clone* 9 (Sf9) suspension cells, which were maintained in SFM II 900 medium (Thermo Fisher Scientific, Waltham, MA, USA) at 27°C and 120 rpm. The ZIKV ZKC2 2016 strain was cultivated in C6/36 cells (MEM with 10% FBS) at 27°C.

The anti-flavivirus E monoclonal antibody (mAb) D1-4G2-4-15 was purchased from Millipore (Billerica, MA, USA). The rabbit anti-ZIKV E polyclonal antibody was purchased from GeneTex (Alton PkwyIrvine, CA, USA). Fluorescein isothiocyanate (FITC)-conjugated goat anti-mouse IgG, horseradish peroxidase (HRP)-conjugated goat anti-rabbit IgG and HRP-conjugated goat anti-mouse IgG were purchased from Abcam (Cambridge, MA, USA). Purified ZIKV-E protein from *Escherichia coli* was purchased from MyBioSource (San Diego, CA, USA).

### Construction of recombinant viruses

Several gene fragments containing prM-E of ZIKV (accession: KX601168.1) were constructed (strategies shown in Fig 1A). The gene fragment “ZI-△-PA” was obtained by overlap PCR, in which the sequences from the N-terminus to the C-terminus were prM-E of ZIKV (with deletion of the stem-transmembrane (ST-TM) region), a linker and protein anchor 3 (PA3, containing three lysin motifs). In ZI-JE-△-PA, the signal peptide of ZI-△-PA was replaced with the signal peptide of Japanese encephalitis virus (JEV). In ZI-GP-△-PA, the signal peptide of ZI-△-PA was replaced with the gp67 signal peptide. Signal peptide replacement was performed by PCR. All of the sequences were optimized for insect cells and synthesized by Sangon Biotech (Shanghai, China).

**Figure 1.**
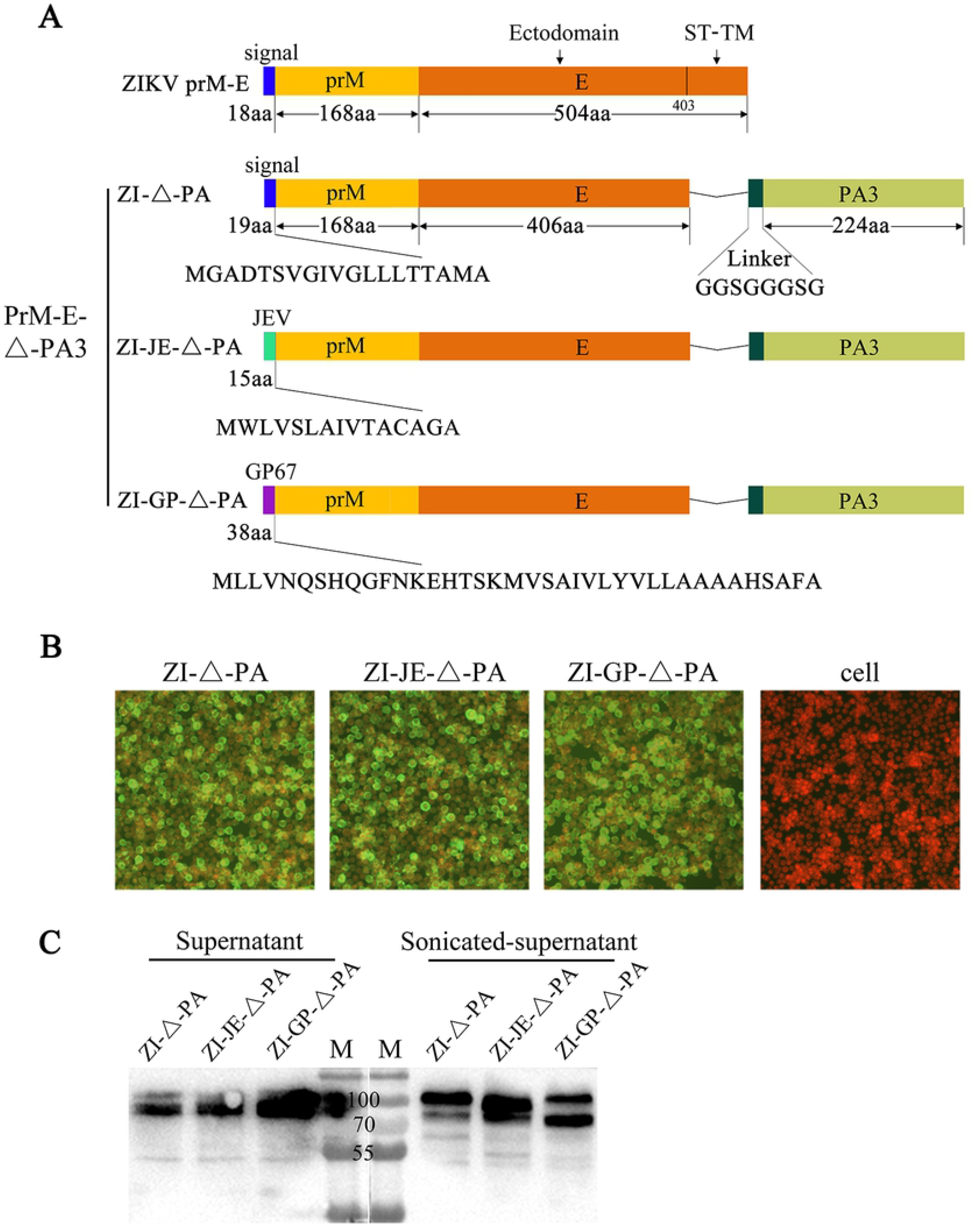
Expression of the fusion protein prM-E- △ -PA3 in the baculovirus expression system. (A) Schematic diagram of the fusion protein prM-E- △ -PA3 fragment. “ △ ” represents a deletion of the stem-transmembrane (ST-TM) region. ZIKV prM-E: the prM-E gene of ZIKV (accession: KX601168.1; does not contain the start codon). ZI-△-PA: the sequences from the N-terminus to the C-terminus were prM-E-△ of ZIKV, a linker and protein anchor 3 (PA3). ZI-JE-△-PA: the signal peptide of ZI-△-PA was replaced with the signal peptide of JEV. ZI-GP-△-PA: the signal peptide of ZI-△-PA was replaced with the gp67 signal peptide. (B) IFA detection of the ZIKV E protein (100×). Recombinant baculovirus-infected Sf9 cells were subjected to IFA for the detection of ZIKV E protein expression. Uninfected Sf9 cells were used as a cell control. (C) WB detection of the ZIKV E protein in baculovirus-infected cells. The proteins were detected in the supernatant or the sonicated supernatant. M represents the protein marker, and the lines in the figure correspond to 100, 70 and 55 kDa.

The above-mentioned target genes were cloned into the baculovirus transfer vector pFastBac Dual under the pH (*Not* I + *Hind* III) and p10 (*Sma* I + *Nhe* I) promoters. Then, the vector was transformed into *E. coli* DH10 Bac/AcMNPV competent cells, and the recombinant baculovirus plasmid (bacmid) carrying two copies of the target genes was obtained. The bacmid was transfected into Sf9 cells to obtain recombinant baculovirus using Cellfectin II Reagent (Invitrogen Co., Carlsbad, CA, USA) according to the manufacturer’s instructions.

### Determination of baculovirus titers

The baculovirus stock titer was measured with a Baculovirus rapid titer kit (TaKaRa, Tokyo, Japan) according to the manufacturer’s instructions. Briefly, samples diluted from 10^-3^ to 10^-5^ were added to Sf9 monolayer cells in a 96-well plate. After incubation at room temperature (RT) for 1 h, the supernatant was replaced with a methyl cellulose overlay. Twenty-seven hours later, the plate was fixed with 4% paraformaldehyde, and blocked with normal goat serum. An *Autographa californica* multiple nuclear polyhedrosis virus (AcMNPV) envelope glycoprotein (gp64) mAb (1:200) was used as the primary antibody, and an HRP-conjugated goat anti-mouse antibody (1:250) was used as the secondary antibody. Blue peroxidase substrate was added and incubated for 3 h at RT. Blue-stained foci in the wells of the highest dilution that contained a reasonable number of foci were counted, and the titer was determined.

### Immunofluorescence analysis (IFA)

For detection of the ZIKV-E protein, Sf9 cells were seeded in 24-well plates and infected with recombinant baculovirus at an MOI of 0.5. Forty-eight hours later, the cells were fixed with 70% cold ethanol and stained with an anti-flavivirus E mAb (1:200) as the primary antibody. Then, a FITC-conjugated anti-mouse antibody (1:200) and Evans blue (1:300) were used. Fluorescence was observed under a fluorescence microscope (Olympus Corp, Tokyo, Japan).

### Western blot (WB) analysis

The recombinant baculovirus was inoculated into Sf9 cells, and the mixture was harvested on the 4^th^ day. The precipitate and supernatant were separated by centrifugation, and the precipitate was sonicated to obtain sonicated supernatant. Samples were denatured in loading buffer at 100°C for 5 min and separated by 10% sodium dodecyl sulfate polyacrylamide gel electrophoresis (SDS-PAGE). Then, the separated proteins were transferred onto nitrocellulose (NC) membranes (GE Healthcare, Little Chalfont, Buckinghamshire, UK) for immunoblot analysis. A polyclonal rabbit anti-ZIKV-E antibody (1:700) was used as the primary antibody, and HRP-conjugated goat anti-rabbit IgG (1:5000) was used as the secondary antibody. SuperSignal West Dual Chemiluminescent substrate (Pierce, Rockford, IL, USA) was added as the chromogen, and bands were imaged using a Tanon-5200 Chemiluminescent Imaging System (Tanon, Shanghai, China).

### GEM preparation

*L. lactis* MG1363 cultured in M17 medium was centrifuged to obtain the bacteria. After washing once with sterile distilled PBS, the bacteria were boiled for 30 min with 10% trichloroacetic acid (TCA) to obtain GEM particles. After washing three times with PBS, the number of GEM particles was counted, diluted to one unit (1 U, 2.5×10^9^ particles per 1U) and stored at −70°C.

### Binding of the fusion protein to GEM particles

Sf9 cells were inoculated with recombinant baculovirus at an MOI of 0.5, and then harvested 4 days later. The supernatant was obtained by centrifugation, and the fusion protein in the supernatant was mixed with GEM at RT for 1 h. After centrifugation at 6000 g/min for 10 min, the precipitate was washed 5 times with PBS and then resuspended in PBS to obtain fusion-GEM complexes.

### TEM analysis and IFA of fusion-GEM complexes

For transmission electron microscopy (TEM) observation, the fusion-GEM complexes were centrifuged to obtain the precipitate. After fixation with 2% glutaraldehyde (no suspension) overnight at 4°C, the precipitate was sliced into ultrathin sections. Then, the samples were stained with uranium acetate and lead citrate, and their features were observed under a transmission electron microscope.

For the IFA, the precipitate from centrifuged fusion-GEM complexes was suspended and blocked with 5% nonfat milk. Then, the cells were stained with an anti-flavivirus E mAb (1:200) as the primary antibody and with FITC anti-mouse IgG (1:200) as the secondary antibody. The stained complexes were applied to slides and observed under a fluorescence microscope (Olympus Corp, Tokyo, Japan).

### Vaccine preparation and immunization

Fusion-GEM complexes (10 μg per mouse) were mixed with a complex adjuvant of ISA 201 VG (Seppic, Paris, France) and poly(I:C) (Sigma, St. Louis, MN, USA). Each milliliter of vaccine contained 350 μl of fusion-GEM complexes, 100 μl of 2 mg/ml poly(I:C) and 550 μl of ISA 201 VG. Female BALB/c mice (6-8 weeks) were randomly divided into 4 groups (n=18/group) and vaccinated intramuscularly (IM) with the vaccines. Mice that received PBS mixed with adjuvant were used as controls. Nine mice from each group were boosted twice at 2-week intervals. Serum was collected at 0, 4, 6 and 8 weeks post-immunization (wpi) and heat inactivated at 56°C for 30 min.

### Detection of serum antibodies

The IgG antibody titers of serum samples were analyzed by an indirect enzyme-linked immunosorbent assay (ELISA). Purified ZIKV E protein was diluted and added to 96-well microtiter plates (Corning-Costar, Corning, NY, USA) at 0.5 μg/well. After incubation at 4°C overnight and washing with PBST, the plates were blocked with 5% nonfat milk. Then, serum diluted 128-fold with blocking buffer was added, followed by HRP-conjugated goat anti-mouse IgG (H+L) (1:2000, BioWorld, St. Louis, MN, USA), IgG1 or IgG2a (1:2000, SouthernBiotech, Birmingham, AL, USA). Tetramethylbenzidine (TMB) was added, and 2 M H_2_SO_4_ was used to stop color development. Absorbance was read at an optical density (OD) of 450 nm using a microplate reader (Thermo Fisher Scientific, Waltham, MA, USA). The results are displayed as the positive index (ratio of sample OD_450_ to control OD_450_).

A virus neutralization assay (VNA) of ZIKV was performed as follows. The serum was serially diluted twofold and mixed with an equal volume of virus (500 PFU/ml) at 37°C for 1 h. Then, 200 μl of the serum-virus mixture was transferred to a monolayer of baby hamster kidney (BHK) cells. After culturing for 1 h, the mixture was replaced with a 2% agarose overlay. The cells were cultured for another 4 days at 37°C. After fixation with paraformaldehyde and staining with crystal violet, a standard 50% plaque reduction neutralization test (PRNT_50_) was performed to calculate the titer.

### B cell and DC activation analysis early after immunization

3, 6 and 9 days after the first immunization, the inguinal lymph nodes of 3 mice in each group were collected and disrupted with the plunger of a syringe. The cell suspensions were stained with anti-CD11c (PE-Cy7), anti-MHCII (PE), anti-CD19 (APC) and anti-CD40 (FITC) antibodies (BD Biosciences, Franklin, CA, USA). Data were acquired on a FACSCalibur flow cytometer (BD Biosciences, Franklin, CA, USA).

### Splenocyte proliferation assay

One week after the last immunization, the spleens of 3 mice in each group were collected and disrupted with the plunger of a syringe. After straining through a 100 μm mesh, red blood cells were lysed in erythrocyte lysis buffer (Solarbio, Beijing, China). Splenocytes (2.5×10^5^) were plated together with the purified ZIKV-E antigen (10 μg/ml) in 96-well plates. After incubation for 44 h, 10 μl/well CCK-8 (KeyGEN Biotech, Nanjing, China) was added to the cells and cultivated for another 4 h. Then, the absorbance was measured at 450 nm using a microplate reader (Thermo Fisher Scientific, Waltham, MA, USA). The proliferation index (PI) was calculated as: (OD_stimulated cultures_ – OD_non-stimulated cultures_)/(OD_non-stimulated cultures_ – OD_control cultures_).

### ELISpot assay and cytokine measurement

A total of 5×10^5^ splenocytes were cultured together with the purified ZIKV-E antigen (10 μg/ml) in 96-well plates. For the ELISpot assay, cells were cultured for 24 h and detected using a commercial kit (MABTECH, Nacka, Sweden) according to the manufacturer’s instructions. For cytokine measurement, cells were cultured for 72 h, and the supernatant was collected for detection using a Meso Scale Discovery (MSD) kit (Meso Scale Diagnostics, Rockville, MD, USA).

### Flow cytometry assays of staining of cell-surface molecule staining

A total of 5×10^5^ splenocytes were cultured and stimulated with the purified ZIKV-E protein (10 μg/ml) for 72 h. Then, the cells were collected and stained with anti-CD4 (FITC), anti-CD8 (PE), anti-CD19 (APC), anti-CD69 (PE/Cy7), anti-CD44 (APC), and anti-CD62L (PerCP-Cy5.5) antibodies (BD Biosciences, Franklin, CA, USA). Data were analyzed using a FACSCalibur flow cytometer (BD Biosciences, Franklin, CA, USA).

### Animal ethics statement

All mice were treated in accordance with the Chinese ethical guidelines for the welfare of laboratory animals (GB 14925-2001). The study was approved by the Animal Welfare and Ethics Committee of the Changchun Veterinary Research Institute.

### Statistical analyses

All experiments were repeated three times, and the results are expressed as the mean ± SD. All statistical analyses were performed by Student’s t-test using GraphPad 8.0.2 software. A p value of ≤ 0.05 was considered statistically significant.

## Results

### Construction and expression of the prM-E-△-PA3 fusion protein

Based on the baculovirus expression system, we constructed several prM-E-△-PA3 fragments fusing the ZIKV prM-E-△ gene (ST-TM region deletion) and the gene for the carrier protein PA3, as shown in Fig. 1A. In the construction strategies, signal peptides from different sources, such as JEV and gp67 (a surface glycoprotein of baculovirus), were considered. IFA of the ZIKV-E protein showed positive results, with bright green fluorescence (Fig. 1B). To determine the localization of the target protein, the supernatants and sonicated supernatants of cell cultures were subjected to WB analysis. The results showed that ZI- △ -PA, ZI-JE- △ -PA and ZI-GP- △ -PA were secreted into the supernatant (Fig. 1C). The IFA and WB results implied that the prM-E-△-PA3 protein reacted with an anti-flavivirus mAb with good antigenicity. When ST-TM was not included, the target protein was secreted.

### Identification of fusion-GEM complexes

To prepare GEM, *L. lactis* was boiled with TCA. In this process, the bacteria were killed, and DNA and proteins were degraded. GEM maintained the original shape and size of the bacterial skeleton, with a smooth surface, and showed good binding with fusion proteins (Fig. 2A). The cell culture supernatants (ZI-△-PA, ZI-JE-△-PA and ZI-GP-△-PA) were selected for binding with GEM and verified by TEM. The results showed obvious floccules on the surface of GEM (Fig. 2B).

**Figure 2.**
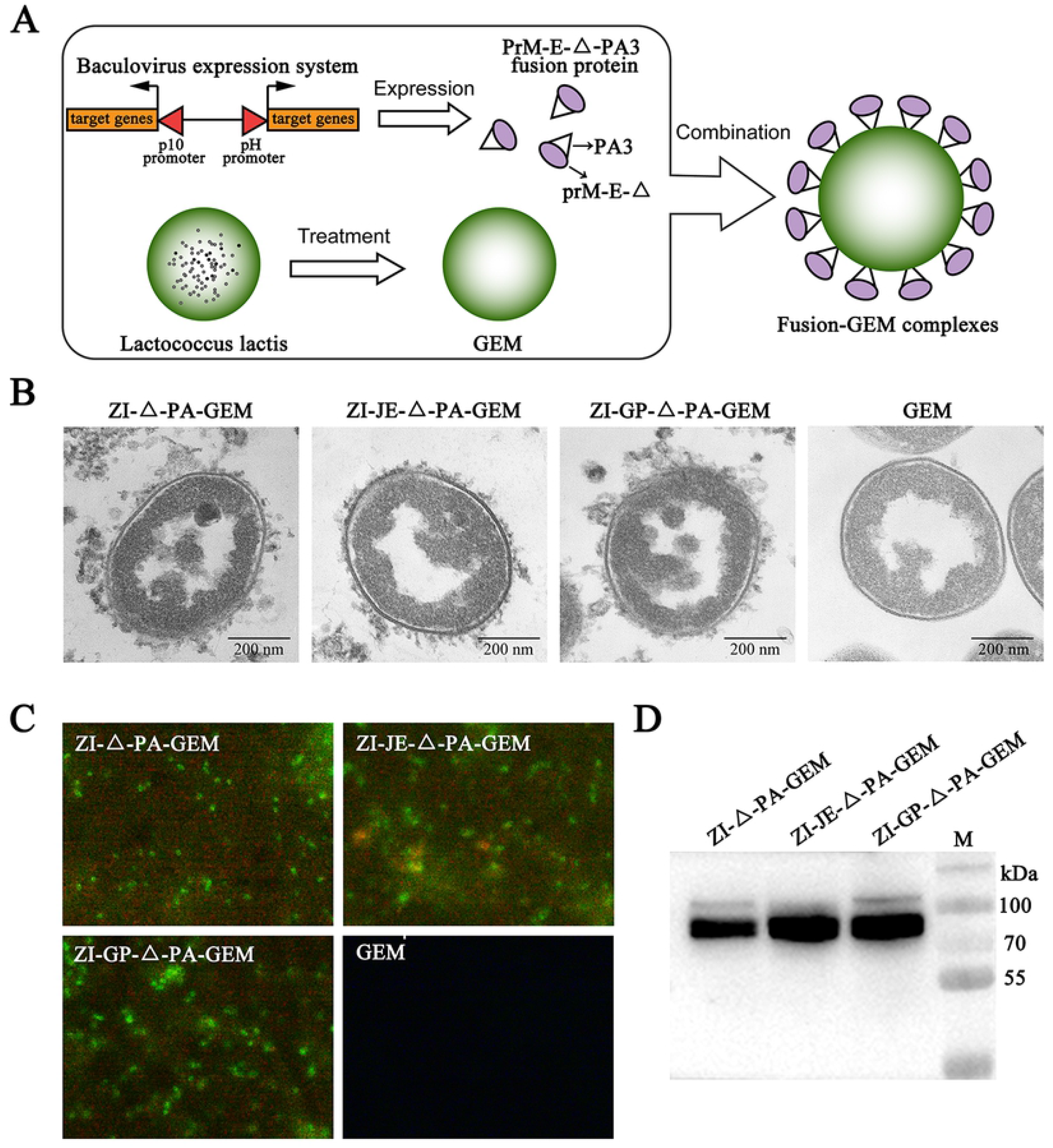
Investigation of fusion-GEM-complexes binding. (A) Combination diagram of the prM-E- △ -PA3 protein binding to GEM. (B) Electron microscopy observation of fusion-GEM-complexes. The scale bars represent 200 nm. GEM was used as the control. (C) IFA detection of ZIKV E protein in fusion-GEM-complexes (1000×). (D) WB detection of the ZIKV E protein in fusion-GEM-complexes. M represents the protein marker.

Furthermore, fusion-GEM complexes with successful binding (ZI-△-PA-GEM, ZI-JE-△-PA-GEM and ZI-GP-△-PA-GEM) were selected for further investigation to verify the ZIKV-E protein in the binding products. The samples showed bright green fluorescence as determined by IFA (Fig. 2C), and the target protein was detected by WB analysis (Fig. 2D). These results indicated that prM-E-△-PA3 with an ST-TM region deletion could successfully bind GEM.

### Fusion-GEM complexes induce ZIKV-specific IgG and neutralizing antibodies

Fusion-GEM complexes (ZI-△-PA-GEM, ZI-JE-△-PA-GEM and ZI-GP-△-PA-GEM) were chosen for immune evaluation in BALB/c mice (Fig. 3A). ZIKV E-protein-specific IgG could be detected in the serum for at least 8 weeks (Fig. 3B). The serum at week 6 was subjected to a VNA, and the results (Fig. 3C) showed that the mean PRNT_50_ titers were all above 10, which is the threshold for protection.

**Figure 3.**
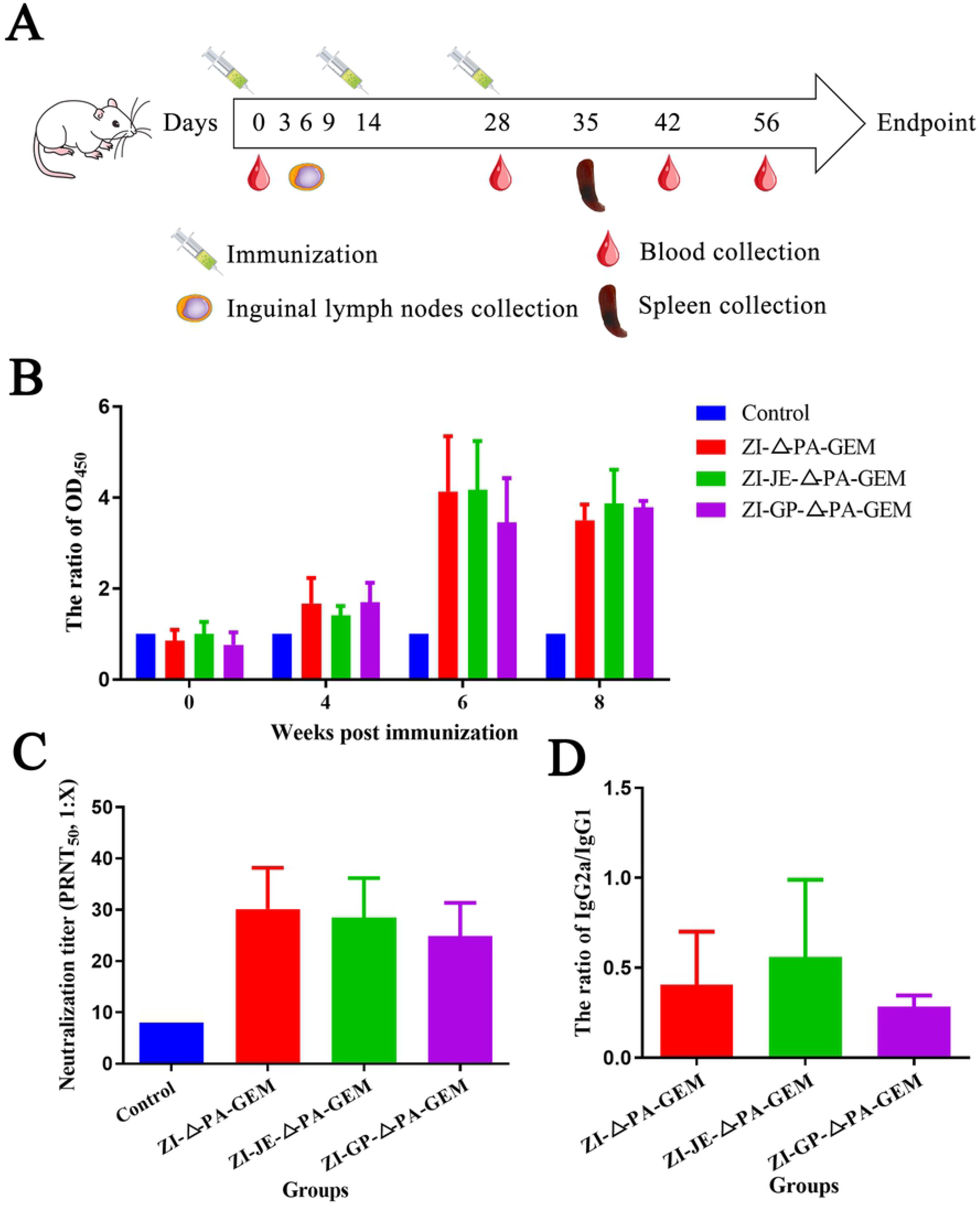
Efficacy of the immunogen in BALB/c mice. (A) Experimental scheme. BALB/c mice (n=18 mice/group) were immunized with ZI-△-PA-GEM, ZI-JE-△-PA-GEM or ZI-GP-△-PA (10 μg) mixed with a complex adjuvant of ISA201 VG and poly(I:C) by intramuscular injection. Mice injected with PBS combined with adjuvant were used as the control. On the 3^rd^, 6^th^ and 9^th^ days after the first immunization, lymphocytes from the inguinal lymph nodes of the mice (3 mice/group/day) were analyzed. Nine mice were boosted twice at 2-week intervals, and sera were collected on days 0, 28, 42, 56 and 70. On the 35^th^ day, splenic lymphocytes from the mice (n=3 mice/group) were analyzed. (B) ZIKV-specific IgG titers in the serum (dilution 128-fold) were measured by ELISA at the indicated time points, displayed as the OD_450_ ratio of sample/control. (C) ZIKV neutralizing antibody titers detected at week 6 by a standard 50% plaque reduction neutralization test (PRNT_50_). (D) The ratios of IgG2a/IgG1 at week 6 were determined by ELISA.

We next examined the quality of this humoral response by IgG2a and IgG1 subisotype-specific ZIKV-E ELISA at week 6 (Fig. 3D). IgG2a/IgG1 ratios lower than 1.0 indicated that ZI-D/ZI-E induced a Th2-biased response.

### B cell and DC activation early after immunization

To investigate the early activation of B cells by the immunogen, mouse lymphocytes from inguinal lymph nodes were analyzed on the 3^rd^, 6^th^ and 9^th^ days after the first immunization. As shown in Fig. 4A, on the 3^rd^ and 6^th^ days after the first immunization, the immune group showed a variable increase in the percentage of CD19^+^ CD40^+^ double-positive cells in the inguinal lymph nodes, with values differing significantly from those of the control group. In addition, ZI-△-PA-GEM caused the strongest activation, with greatest significant differences compared to the control group.

**Figure 4.**
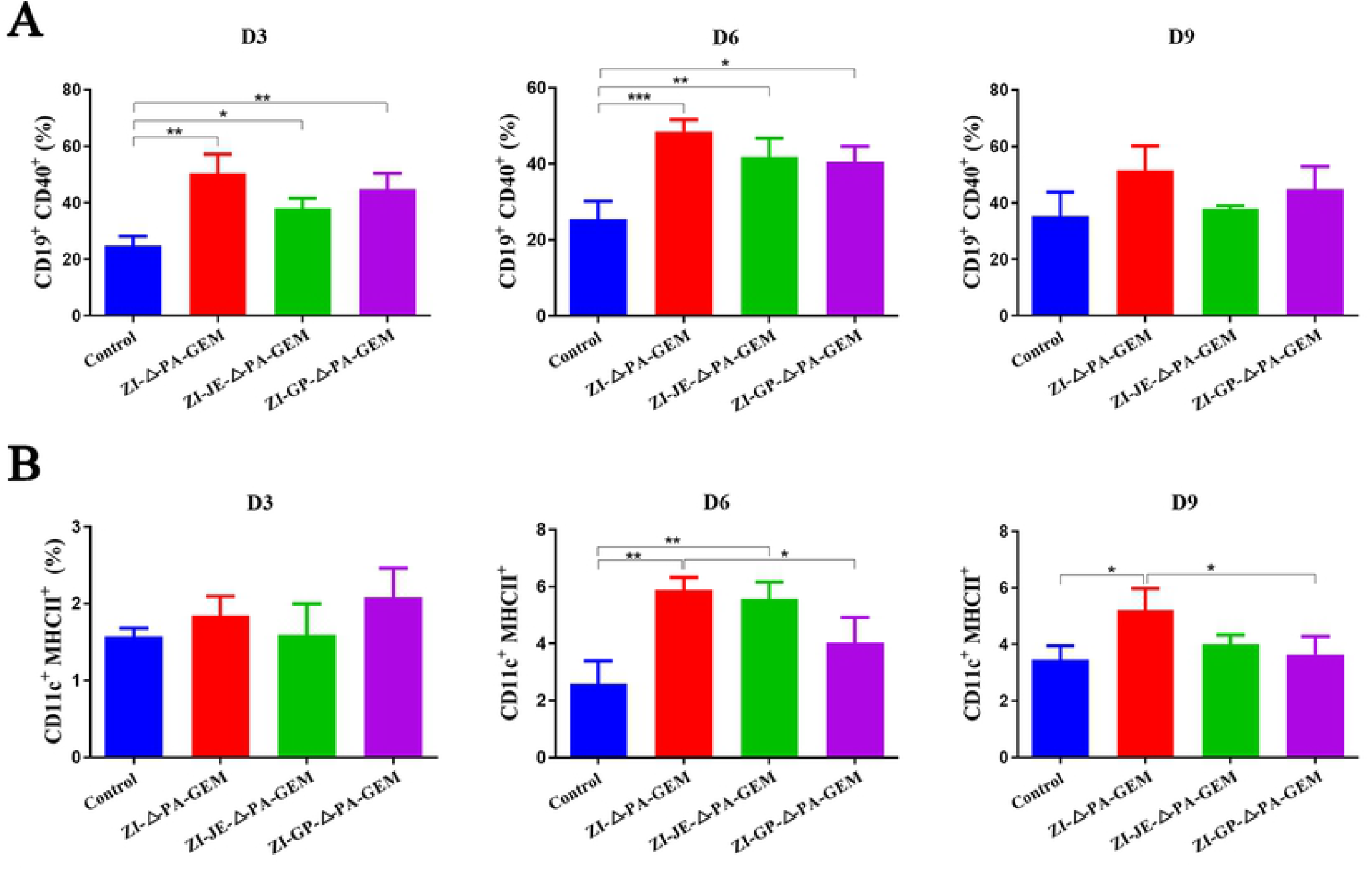
Detection of B cell and DC activation early after immunization. On the 3^rd^, 6^th^ and 9^th^ days after the first immunization, lymphocytes from the inguinal lymph nodes of the mice were analyzed by flow cytometry. (A) B cell activation, displayed as CD19^+^ CD40^+^. (B) DC activation, displayed as CD11c^+^ MHCII^+^. The data are expressed as the mean ±SD for each group. *p<0.05; **p<0.01; ***p<0.001; ****p<0.0001.

Next, the activation of DCs was analyzed by detecting the proportion of CD11c^+^ MHC II^+^ cells (Fig. 4B). On the 3^rd^ day, there was slight activation of DCs, but there was no significant difference among the groups. On the 6^th^ and 9^th^ days after the first immunization, the immune groups had an increased percentage of CD11c^+^ MHC II^+^ double-positive cells. On the 9^th^ day, the activation levels of some groups decreased, but the ZI- △ -PA-GEM group still showed significant differences from the control group.

### Splenocyte proliferation after ex vivo restimulation

One week after the last immunization, the effect of the immunogen on splenocyte proliferation responses was evaluated. After ZIKV-E protein stimulation, splenocytes from immunized mice proliferated more efficiently than those of the control mice (Fig. 5A). Moreover, the ZI-△-PA-GEM group showed the greatest increase in proliferation.

**Figure 5.**
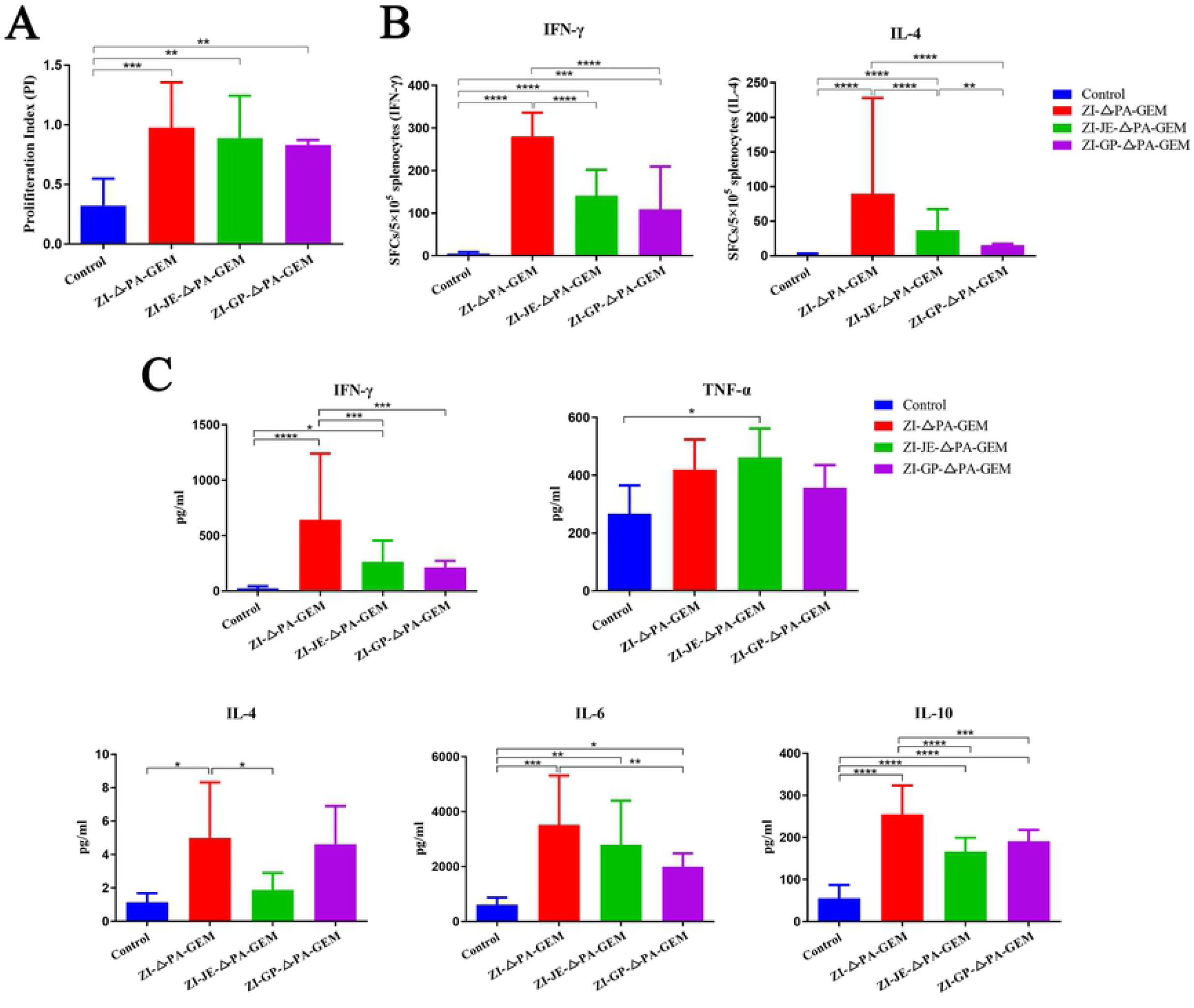
Splenocyte proliferation analysis. On the 35^th^ day after immunization, splenic lymphocytes from the mice were harvested and restimulated with purified ZIKV E protein. (A) After *ex vivo* culture for 44 h, the proliferative index was determined using a CCK-8 assay. (B) After culture for 24 h, the levels of IFN-γ and IL-4 secreted by the splenocytes were quantified by the ELISpot assay. (C) After culture for 72 h, the cytokine levels in splenocyte culture supernatants were measured by MesoScale Discovery (MSD). The data are expressed as the mean ±SD for each group. *p<0.05; **p<0.01; ***p<0.001; ****p<0.0001.

Next, the ability of splenocytes to produce IFN-γ and interleukin-4 (IL-4) in response to specific antigens in vitro was evaluated. Through an enzyme-linked immunosorbent spot (ELISPOT) assay, significantly stronger IFN-γ and IL-4 responses were detected in the immune groups (Fig. 5B). Next, we tested the levels of multiple cytokines in the supernatants of stimulated splenocytes from all groups. The levels of the cytokines Th1 (IFN-γ) and Th2 (IL-4, IL-6 and IL-10) were significantly higher in the ZI-△-PA-GEM group than in the control group (Fig. 5C). The tumor necrosis factor (TNF)-α levels were also increased slightly, but the difference was not significant. In the signal peptide replacement groups (ZI-JE-△-PA-GEM, ZI-GP-△-PA-GEM), the levels of the above cytokines were also increased to varying degrees. Moreover, the trend of IFN-γ and IL-4 cytokine levels in the splenocyte culture supernatants was consistent with the ELISPOT results. These data suggested that ZI-△-PA-GEM could induce immune cell proliferation and that both Th1 and Th2 responses were elicited.

### Lymphocyte activation assays

To analyze the activation of CD4^+^ T, CD8^+^ T and B cells after immunization, we assessed the proportion of CD69^+^ splenic lymphocytes. CD69 is one of the earliest lymphocyte surface markers detected after antigen stimulation [18]. The proportions of CD4^+^ CD69^+^ T, CD8^+^ CD69^+^ T and CD19^+^ CD69^+^ B cells in the ZI-△-PA-GEM group were significantly higher than those in the control group (Fig. 6A). This result indicated that ZI-△-PA-GEM with a complex adjuvant of ISA201 VG and poly(I:C) promoted T/B cell activation.

**Figure 6.**
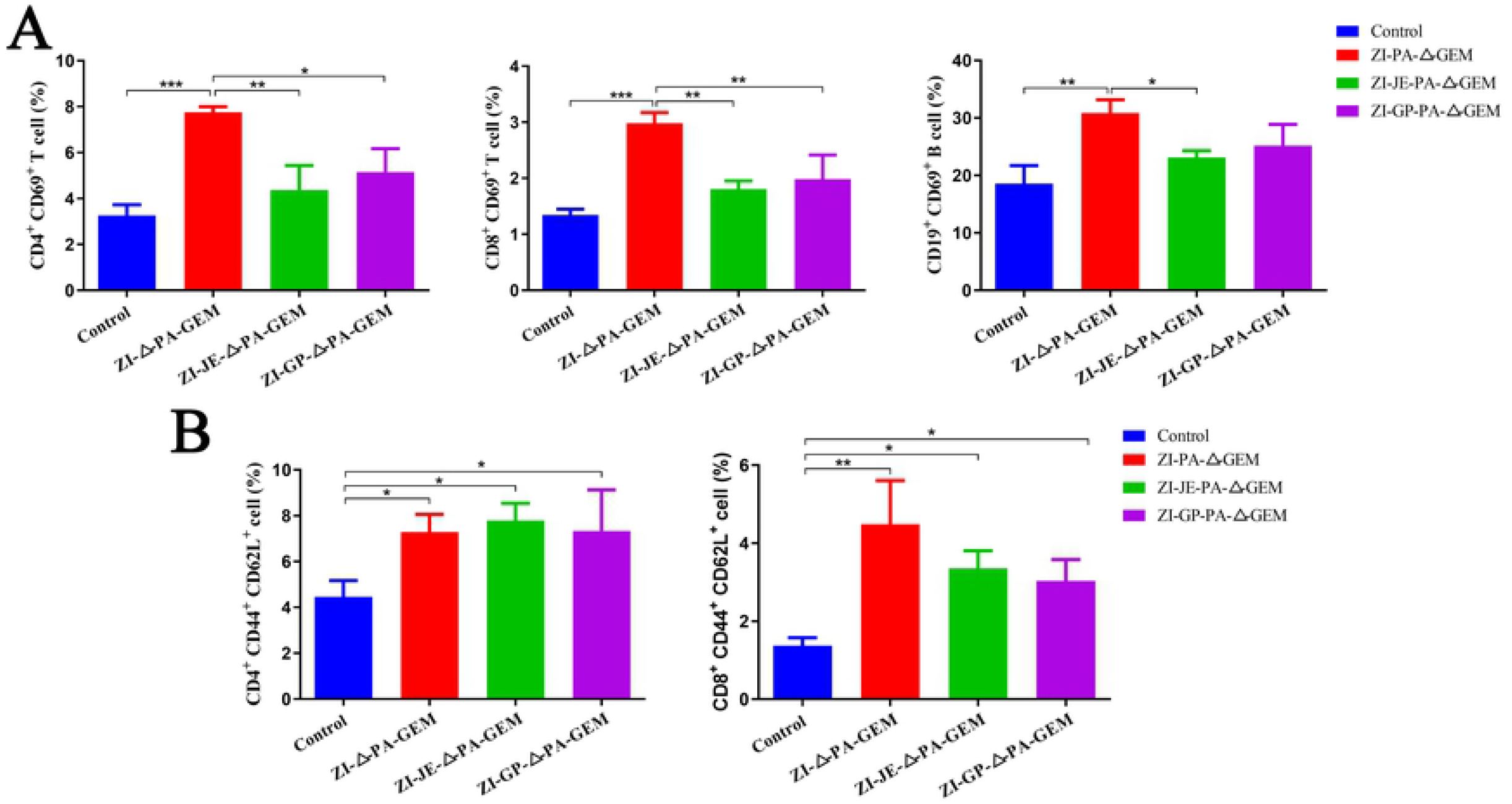
Lymphocyte activation analysis. On the 35^th^ day after immunization, splenic lymphocytes from the mice were restimulated with purified ZIKV E protein and cultured *ex vivo* for 72 h. (A) The activation of CD4 T cells (CD4^+^ CD69^+^), CD8 T cells (CD8^+^ CD69^+^) and B cells (CD19^+^ CD69^+^) was measured by flow cytometry. (B) The proportion of central memory T cells (TCMs) among CD4^+^ or CD8^+^ T cells was evaluated by the presence of CD44 and CD62L surface markers. The data are expressed as the mean ±SD for each group. *p<0.05; **p<0.01; ***p<0.001; ****p<0.0001.

Next, we detected the proportion of central memory T cells (TCMs; CD44^+^ CD62L^+^) among splenic lymphocytes. As shown in Fig. 6B, the proportion of TCMs among CD4^+^ and CD8^+^ T cells was significantly increased by ZI-△-PA-GEM, ZI-JE-△-PA-GEM and ZI-GP-△-PA-GEM.

## Discussion

The GEM-PA display system was first reported by Steen et al [19], who combined the peptidoglycan skeleton of heat-treated *L. lactis* with a foreign protein fused to the PA domain. GEM is a peptidoglycan skeleton of the cell wall obtained by the acid boiling of gram-positive bacteria (food-grade non-gene-modified *L. lactis*). PA is the C-terminal domain of the AcmA protein (an important peptidoglycan hydrolase in *L. lactis*). PA contains a lysin motif (LysM), which can noncovalently bind to peptidoglycan, thus anchoring to the cell wall of GEM [20]. GEM and PA are the vectors and carrier proteins of the system, respectively. Target proteins fused to PA can be anchored on the surface of GEM, namely GEM surface display system. Due to several advantages, this system has been studied and applied in many vaccine studies [16, 21, 22]. Natural PA contains three LysMs, and serine, threonine and aspartic acid are used as the interval sequences between the repeat sequences [23]. Multiple LysMs might form multivalent domains, thus increasing the anchoring activity of PA, but the anchoring activity of PA containing four LysMs was significantly reduced [20, 23]. Based on our previous research [24], we chose PA3 (containing three LysMs) as the carrier protein and an 8-amino acid sequence as the linker.

In the development of ZIKV DNA vaccines, Dowd [25] exchanged the signal sequence of prM-E with the analogous region of JEV to improve expression. The ST-TM was also exchanged with the corresponding JEV sequences to improve virus-like subviral particle secretion. In this study, we replaced the signal peptide of ZIKV with the corresponding sequence of JEV or gp67. In addition, we initially replaced the ST-TM region of ZIKV with the corresponding JEV region (data not shown). After fusion of prM-E with PA3, the target protein was hardly secreted into the supernatant and could be detected in only the sonicated supernatant. Moreover, the fusion protein with the ST-TM region could not bind to the surface of GEM (as detected by TEM, showing a smooth GEM surface). The ST-TM region likely affected the correct expression or function of the PA3 anchor protein. Therefore, we further deleted the ST-TM region, and the fusion protein was secreted into the supernatant and successfully bound to the surface of GEM. In ZIKV DNA and virus-like particle (VLP) vaccine construction strategies, the ST-TM region is beneficial for protein expression and packaging [25–27]. However, in this study, when the target protein was coexpressed with exogenous protein PA3, deletion of the ST-TM region was beneficial for the binding of the target protein to GEM.

ZI-△-PA-GEM stimulated the body to produce specific antibodies against the ZIKV-E protein for at least 8 weeks. The PRNT_50_ value of NAb was above 1:10. There is only one serotype of ZIKV [28], and NAb titers >10 have been found to correlate with protective efficacy [26, 29, 30]. Our results indicated that ZI-△-PA-GEM could induce potent E-specific IgG antibodies, as well as NAb responses. In addition, the antibody subtype was Th2 biased, which might be advantageous for protection in some ways and has been reported for vaccines [24, 31–34].

As a surface molecule of B cells, CD40 plays an important role in promoting the differentiation of B cells into regulatory cells [35]. The coreceptor CD19 plays an essential role in mediating the spreading of B cells and enhancing signaling through the B cell receptor (BCR) in response to membrane-bound antigen [36]. In this study, in the early stage after the first immunization, ZI- △ -PA-GEM stimulated B cell activation, with an increase in CD19^+^ CD40^+^ double-positive cells. Activated B cells can produce plasma cells that generate high-affinity antibodies and memory cells, thus protecting the body against infectious diseases [37]. Moreover, ZI- △ -PA-GEM recruited and/or activated DCs in secondary lymphoid organs and significantly stimulate the expression of MHC II molecules, with an increase in CD11c^+^ MHC II^+^ double-positive cells. CD11c is a sign of DC maturation, and DCs are the most professional antigen-presenting cells (APCs) and the only APCs that can stimulate the initial T cell response [38]. In addition, DCs are an important factor regulating the differentiation of Th0 cells into Th1 or Th2 cells. MHC II molecules are important for presenting antigens on the surface of DCs [39, 40], mainly exogenous antigens, and play multiple roles in inducing protective immunity after vaccination [41]. The presentation of antigen peptides on the MHC II molecules of DCs can induce initial CD4^+^ T cell activation [39].

Functional regulation of Th subsets and the cytokine environment is beneficial to protective immunity [42]. ZI- △ -PA-GEM induced significant cytokine secretion, including IFN-γ, IL-4, IL-6 and IL-10. Th1 cells produce important cytokines for the production of cytotoxic T cells and complement-fixing antibodies. Th2 cells produce a series of cytokines, that support antibody production to protect against extracellular pathogens [43]. The cytokines induced by vaccines are synergistic and rarely work alone, which complicates the interpretation of immune correlations [44]. The differential cytokine levels between the immunized and control groups may contribute to protection against flaviviruses [45]. Moreover, cytokines affect the differentiation of memory T cells [42]. There are two subpopulations of memory cells: effector memory T cells (TEMs; migrate mainly to peripheral tissues) and TCMs (localize to the lymph nodes, blood and spleen) [42]. Here, the proportion of TCMs among splenic lymphocytes stimulated by ZI- △ -PA-GEM was significantly higher, which was beneficial for rapid stimulation of the immune response during antigen restimulation.

High levels of T cell activation were also detected, especially with ZI-△-PA-GEM. Although there was no clear final conclusion concerning whether the cell-mediated immune responses against ZIKV contributed to protection, the results of a previous study support that T cells play an important role in protecting against ZIKV [4, 46]. CD8^+^ T cells limit the spread of viruses by recognizing and killing infected cells or secreting specific antiviral cytokines, and CD4^+^ T cells contribute to cytokine production and support the generation and maintenance of antibody and CD8^+^ T cell responses [45].

Potent B cell/DC activation, cytokine responses, specific splenocytes capable of proliferation, and T cell responses, together with the production of specific antibodies, are likely to jointly contribute to protection in mice. Taken together, our data indicate that ZI-△-PA-GEM is capable of inducing humoral and cellular immune responses in mice that may protect against ZIKV infection.

## Author Contributions

**Conceptualization:** Hualei Wang, Xianzhu Xia, Hongli Jin

**Data curation:** Hongli Jin, Yujie Bai, Hualei Wang

**Formal analysis:** Cuicui Jiao, Jianzhong Wang

**Funding acquisition:** Hualei Wang

**Investigation:** Pei Huang, Di Liu, Yumeng Song

**Methodology:** Hongli Jin, Yujie Bai, Cuicui Jiao, Mengyao Zhang, Nan Li

**Project administration:** Hongli Jin, Xianzhu Xia, Hualei Wang

**Resources:** Na Feng, Yongkun Zhao, Tiecheng Wang, Yuwei Gao

**Software:** Hongli Jin, Zhiyuan Gong, Entao Li, Shengnan Xu

**Supervision:** Yuwei Gao, Songtao Yang.

**Validation:** Hualei Wang, Xianzhu Xia.

**Visualization:** Hongli Jin, Yongkun Zhao, Tiecheng Wang.

**Writing – original draft:** Hongli Jin.

**Writing – review & editing:** Hongli Jin, Songtao Yang, Hualei Wang

## Acknowledgments

We thank Dr. Xuejin Su for helping us collect and analyze the flow data. We also thank Jingbo Huang, Jingxuan Sun, Xingqi Liu and Jiaxin Dai for kindly help in animal experiments.

## Competing Interests

The authors have declared that no competing interests exist.

## Notes

### Competing Interest Statement

The authors have declared no competing interest.

